# Chromium/cadmium plays a pivotal role to emerge amoxicillin resistant *Staphylococcus aureus*

**DOI:** 10.1101/2023.02.28.530213

**Authors:** Tajreen Naziba Islam, Foujia Samsad Meem, Rahena Yasmin, Mohammed Badrul Amin, Tania Rahman, David H. Dockrell, Md Mohasin

## Abstract

**Rationale:** The rapid emergence of resistant bacteria is occurring worldwide, endangering the efficacy of antimicrobials. Apart from horizontal gene transfer and plasmid mediated antimicrobial resistance (AMR) acquisition, co-exposure of heavy metals and antibiotics cause to emerge AMR Enterobacteriaceae. Heavy metals and antimicrobials co-exist in many environmental settings. We hypothesized that heavy metals and lower dose of antibiotic co-exposure may alter levels of antimicrobial susceptibility and facilitate to emerge AMR bacteria.

**Methods:** The growth kinetics of antimicrobial susceptible Staphylococcus aureus ST80 was carried out in the presence of chromium/cadmium salt and a lower dose of antibiotics. Subsequently, the antimicrobials susceptibility patterns of heavy metals pre-exposed for 48 hours Staphylococcus aureus ST 80 was determined by Kirby-Bauer disc diffusion method.

**Results:** The antimicrobial susceptibility profile revealed that the zone of inhibition (ZOI) for ampicillin, amoxicillin, ciprofloxacin and doxycycline significantly decreased in chromium pre-exposed Staphylococcus compared to unexposed bacteria. However, cadmium pre-exposed bacteria only showed significant decreased ZOI for amoxicillin. Moreover, the MIC of amoxicillin was increased by 8-fold in chromium and 32-fold in cadmium with a low-dose of amoxicillin co-exposed bacteria. Besides, the RT-qPCR data demonstrated that chromium and a low-dose of amoxicillin pre-exposed significantly increased the mRNA expression of femX (25-fold), mepA (19-fold) and norA (17-fold) in S. aureus.

In essence, minimum levels of chromium/cadmium and a MIC of amoxicillin exposure induced efflux pumps, which might responsible to emerge amoxicillin resistant S. aureus.

## 1. Introduction

Antibiotic resistance is one of the major global problems and threatens the usefulness of nearly all antibiotics that was discovered to alleviate microbial infections (1). Several studies have proved that antimicrobial agents other than antibiotics have the capability to cause antibiotic resistance through a co-selection process (2, 3). In addition to antibiotic resistance, heavy metal contamination is another severe ecological problem (4, 5). Antibiotic and heavy metals co-exist in the environment such as in the gastrointestinal tract, animal manure, and poultry farming sites (6–9). For instance, arsenic and antibiotics are widely used in poultry farms as growth promotion and disease control agents thus the gut microbiota of domestic animals is getting exposed to both antibiotics and heavy metals. Additionally, fertilizers made from manure and sewage sludge containing those substances are extensively used in the agricultural soil that eventually leach into the water (3, 10). As a result, humans are exposed to these heavy metals through the contamination of the food chain as heavy metals are not easily bio-degradable (4, 11). Several studies have reported that the elevated concentration of heavy metals can induce antibiotic resistance (3, 7, 12). According to Seiler & Berendonk, 2012, the combined effect of heavy metals or metals and antibiotics discharged into soil and water bodies maybe responsible for the spread of antimicrobial resistance as well as the evolution of multidrug resistance.

Moreover, Peltier et al. reported that co-exposure of metal like Zinc and oxytetracycline promotes microbial resistance towards oxytetracycline (13). Similarly, another study demonstrated that addition of Copper (Cu) in agricultural soils not only arises Cu resistance but also co-selects for resistance to ampicillin, chloramphenicol and tetracycline (14). Recently, Chen et al., has demonstrated that the growth of LSJC7, an Enterobacteriaceae strain significantly increased in arsenate and tetracycline co-exposure milieu compared with only tetracycline treated growth (9). The possible mechanism of such condition could be due to the presence of heavy metals induced growth of microbial community having resistance genes beforehand, or through co-selection process heavy metals and antibiotics together stimulated microbial resistance to the antibiotics that was previously sensitive (9). In this study, we hypothesized that bacterial growth with a lower dose of heavy metals or antibiotics may alter antimicrobials susceptibility and the expression patterns of efflux pumps genes, which are responsible to emerge antibiotic resistant bacteria. The antimicrobial sensitivity profile and growth kinetics of *Staphylococcus aureus* ST80 was determined in the presence of chromium or cadmium salts. This study has revealed that growth of *S. aureus* with a lower dose of chromium or cadmium and amoxicillin was significantly increased the minimum inhibitory concentration (MIC) of amoxicillin and also facilitate to acquire amoxicillin resistance through alter the expression of efflux pumps and femX gene.

## 2. Methods and Materials

### Place of Study

This study was carried out at the Infection and Immunity Laboratory in the Department of Biochemistry and Molecular Biology, University of Dhaka, Bangladesh.

### Collection of *Staphylococcus aureus* ST80

Several skin, soft-tissue, respiratory, bone, joint, and endovascular disorders are associated with *Staphylococcus aureus*, which is responsible for numerous infectious diseases (15). In this study, the *Staphylococcus aureus* ST80 was isolated from processed raw meat (meat ball) of a local restaurant by Food Microbiology Laboratory, Laboratory Sciences and Services Division, International Center for Diarrhoeal Disease Research, Bangladesh (ICDDR,B).

### Measurement of Heavy metal levels in Industrial wastewater

Samples of Industrial wastewater were collected from Buriganga and Dhaleswari river. To compare the effectiveness of effluent treatment process (ETP), we had also collected wastewater directly from some tanneries and textile industries before ETP and after ETP. Briefly, to determine the levels of heavy metals wastewater was dissolved in 65% nitric acid (HNO_3_) in order to minimize precipitation by bringing the pH lower than 2.0 (Hassan et al., 2015). 5mL of 65% concentrated HNO_3_ was added in each volumetric flasks containing 100 mL of wastewater. It was then gently boiled until complete dissolved on a hot plate in a fume hood, cooled prior to filtration using Whatman™ qualitative filter paper (16). Finally, each water samples were loaded onto Flame Atomic Absorption Spectrophotometer for analysis and detection of heavy metals like Cr, Cd, and Pb. Before running the samples in AAS, the instrument was first calibrated with chemical standard solutions according to the manufacturer’s instructions (17).

### Preparation of Agar Media for Culturing the Bacteria

To culture bacteria in a solid media, Tryptone Soya Agar (TSA) media was used. The TSA powder was measured using a balance (Shimadzu ELB200, Japan) and taken into a conical flask. After adding desired volume of distilled water in the conical flask, the mixture was mixed with the help of a magnetic stirrer. It was then autoclaved at 121°C under 15 psi for 20 minutes. The media preparation was carried out in the biosafety cabinet in order to avoid contamination. The media was poured in the petridish and allowed to solidify for a few minutes. The *Staphylococcus aureus* ST80 was streaked on the prepared agar plate from the collected culture plate using the inoculation loop. The plate was then placed in the incubator (Memmert, Germany) at 37°C and allowed to grow the bacteria overnight. After 16 hours, the plate containing the colony of the bacteria was stored in the refrigerator at 4°C.

### Dose-response growth kinetics in presence of chromium (Cr^6+^) salt

*Staphylococcus aureus* ST80 was grown in 0.5mM, 1mM, 3mM, 5mM, 10mM, 50mM, 100mM chromium salt in Tryptone Soya broth (TSB) media and O.D value was recorded at 600nm using UV-Vis spectrophotometer (Thermo-Scientific).

### Dose-response growth kinetics in presence of cadmium (Cd^2+^) salt

*Staphylococcus aureus* ST80 was grown in 0.005mM, 0.01mM, 0.025mM, 0.05mM, 0.075mM, 0.1mM, 0.3mM, 0.5mM, 0.75mM cadmium salt in Tryptone Soya broth (TSB) media and O.D value was recorded at 600nm using UV-Vis spectrophotometer.

### Standardization of Tryptone Soya Medium

TSB medium containing bacterial solution was carried out for the serial dilution using 0.9% saline under the laminar flow. 1mL TSB media was transferred in a tube, centrifuged at 4000 rcf for 3 minutes. The liquid suspension was discarded, and the bacterial pellet was diluted with 1mL saline solution. This step was repeated two times to achieve pure bacterial culture. These media were then standardized through spectrophotometry method. 1.0×10^8^ CFU/ml of bacterial concentration were assured in each of the cultured sample in TSB medium, which was represented by 0.125 OD.

### Pre-screening Antibiotic susceptibility pattern in presence of chromium and cadmium (for Table)

Bacteria was grown in three conical flasks with concentration of 0.0 mM, 0.5 mM, 1.0 mM chromium containing TSB media. For cadmium, 2 mM stock solution of salt (CdCl_2_.H_2_O) was prepared by dissolving 0.02g CdCl_2_.H_2_O (Cd^2+^) in 50 mL distilled water and heated and mixed using magnetic stirrer to dissolve the solvent completely. Then concentration of 0.0 mM, 0.05 mM, 0.1 mM cadmium containing TSB media were prepared from stock solution in another three conical flask. 10μL equivalent to 1.0×10^8^ cfu/mL of standardized bacterial suspension was added in each flask. The liquid broth flasks were then placed into shaking incubator at 37°C, 180 rpm for 12 hours. Then 100μl of pre-exposed bacterial solution from each conical flask was spread on several Mueller-Hinton agar plates using sterile spreader. Antibiotic discs were then impregnated on the surface of agar plate using sterile forceps. 18 agar plates were prepared and out of 9 antibiotics 3 antibiotics were placed in each agar plate. All the plates were then incubated at 37°C for a period of 16-24 hours. The diameter of the zone of inhibition around the disc was measured using a millimeter scale and compared to the CLSI reference table to determine if the organism is susceptible, intermediate or resistant against the antibiotic agents tested.

### Metal and Antibiotic Analysis on pre-exposed Bacterium

To determine the effect of Cr^6+^ and Amoxycillin on *Staphylococcus aureus* ST80 growth, the bacterium was grown on different conditions.

a. Cr^6+^ pre-exposed SA
b. Cr^6+^ exposed in Cr^6+^ pre-exposed SA
c. Amoxycillin exposed in Cr^6+^ pre-exposed SA
d. Amoxycillin and Cr^6+^ co-exposed in Cr^6+^ pre-exposed SA
e. Amoxycillin exposed in Cr^6+^ unexposed SA
f. Amoxycillin and Cr^6+^ co-exposed in Cr^6+^ unexposed SA

### Co-exposure effect of amoxicillin treatment upon bacterial growth in presence of Cr^6+^ salt

1.5g TSB was taken in each of two conical flasks. To prepare 0.5mM chromium containing liquid broth, 0.0074g of K_2_Cr_2_O_7_ was measured and added into one conical flask. Then the reagents were dissolved using 50mL distilled water and sterilized by autoclaving the media. 50mL of each media was then poured into two different centrifuge tubes. 0.06 μg/mL of Amoxicillin was added into one chromium containing tube and one TSB media containing tube. 10μL equivalent to 1.0×10 8 cfu/mL of standardized bacterial suspension was added in each centrifuge tube except blank tube. The liquid broth tubes were then placed into shaking incubator at 37°C, 180 rpm and the optical density of each tube was taken at 600 nm in every hour (9). Before that, blank of each condition was performed.

### Co-exposure effect of amoxicillin treatment upon bacterial growth in presence of Cd^2+^ salt

In each of two conical flasks, 1.5g TSB was taken. Now to prepare 0.025mM Cd^2+^ containing media; 0.625 mL stock of cadmium salt was added in one conical flask and the final volume was made 50mL with distilled water. All liquid broth media was then autoclaved and allowed to cool down in room temperature. 25mL of each media was poured into four different centrifuge tubes. 0.06 μg/mL of Amoxicillin was added into one cadmium containing tube and one TSB media containing tube. One of each tubes were provided with 10μL of standardized bacterial suspension and placed into shaking incubator at 37°C, 180 rpm and the optical density of each tube was taken at 600 nm in every hour (9). Blank of each condition was performed before measuring optical density of different conditions.

### Determination of Minimum Inhibitory Concentration of SA upon treatment with a Sub-lethal dose of Chromium salt K_2_Cr_2_O_7_ (Cr^6+^) and Amoxicillin Using Agar Dilution Method

SA was grown in Liquid Broth media supplemented with 0.5mM chromium, 0.06μg/ml Amoxicillin and a medium containing 0.5mM chromium and 0.06μg/ml Amoxicillin both respectively for 12 hours, 24 hours and 48 hours in a continuous batch culture system (fresh TSB culture media with or without chromium salt and amoxicillin supplement was changed in 12 hours interval). After 48hours of exposure in stressed condition, the bacterial culture was purified using NaCl and the turbidity was adjusted to O.D_600_ value 0.125. Solutions of 25 mL Tryptone soya Agar (TSA) were prepared for each condition and autoclaved. Before transferring the solution to the plate 0.06μg/mL, 0.125μg/mL, 0.25μg/mL, 0.5μg/mL concentration of antibiotics were added into the conical flasks (Yao et al., 2019). The control media contained no antibiotic. The antibiotics were added after cooling down the medium to 50°C cause higher temperature may inactive the antibiotic and in low temperature the agar will begin forming solid clumps. The media was poured into petri dishes and solidified. After that, 10μL of bacterial culture from each condition (equivalent to 10^5^cfu) was spotted (total three identical spot) in each agar plate. The plates were placed in incubator for 16 hours at 37°C and the MIC of each condition was observed (18). The MIC of this bacterium (grown in presence of both chromium salt and amoxicillin) increased 8fold compared to control as the bacterial growth was found in the media containing 0.50μg/ml concentrations of Amoxicillin.

### Evaluation of Minimum Inhibitory Concentration Using Agar Dilution Method for CdCl_2_.H_2_O (Cd^2+^)

25mL Tryptone soya Agar (TSA) plates were prepared for each condition. Then the autoclaved 25mL TSA was transferred and 0.06μg/mL, 0.125μg/mL, 0.25μg/mL, 0.5μg/mL, 1.0μg/mL, 2.0μg/mL, 4.0μg/mL concentration of antibiotics were added into the conical flasks (Yao et al., 2019). The control media contained no antibiotic. The antibiotics were added after cooling down the medium to 50oC cause higher temperature may inactive the antibiotic and in low temperature the agar will begin forming solid clumps. The media was poured into Petri dishes and solidified. After that, 10μL of standardized bacterial suspension (equivalent to 105cfu) containing inoculum was spotted (total three identical spot) in each agar plate. Thereafter, the bacterial plates were incubated for overnight culture at 37°C and the MIC of each condition was observed (18). The bacteria were grown in presence of both cadmium and amoxicillin (maximum 2.0 μg/mL), demonstrating 32-fold increase of MIC compared to control. Earlier, *Staphylococcus aureus* containing bacterial suspensions were prepared from Tryptone soya broth culture medium, which was pretreated with or without 0.025 mM Cd^2+^, 0.06μg/mL Amoxicillin or a combination of 0.025 mM Cd^2+^ and 0.06 μg/mL Amoxicillin for 12 hours, 24 hours and 48 hours in a continuous batch culture system.

### Antimicrobial susceptibility test (Disc-Diffusion Method)

To perform the Kirby-Bauer disc diffusion method, Mueller-Hinton Agar medium is best considered according to Clinical Laboratory Standards Institute (CLSI guideline, 2017) for Antimicrobial Susceptibility Test. To determine the difference of zone of inhibition, this experiment was carried out in 0.95g of MHA, measured using a balance (Shimadzu ELB200, Japan) and dissolved in distilled water in order to make 25 mL solution for each 100mm petri dish. All the mixtures were autoclaved at 121°C under 15 psi for 20 minutes. The mixtures were then transferred from the conical flask into petri dishes and allowed to solidify for a few minutes. Bacteria from both the MIC value under chromium and cadmium stressed was grown in TSB media. Then 100μL of respective bacterial suspension was spread over the agar plates using sterilized spreader and the filter paper discs of Amoxicillin were carefully dispensed on the surface of each agar plates using sterile forceps. All these steps were carried out in the laminar flow to maintain aseptic condition. The plates were then incubated at 37°C for 16 hours. Afterwards, the diameter of the zone of inhibition around the disc was measured.

### Reverse transcription qPCR for evaluation the expression patterns of efflux pumps and femx gene

#### RNA Extraction and cDNA synthesis

*S. aureus* is lysozyme resistant due to the presence of modified peptidoglycan layer (19), hence the specialized lysis steps are required for RNA isolation other than lysozyme. Previous studies were evaluated several methods to achieve high quantity and quality RNA (20). To recover maximum yield of RNA, the simple phenol method as the most effective one for cell lysis compared to commercially available RNA extraction Kit (20). Here, we have used Monarch® Total RNA Miniprep Kit to conduct subsequent steps of RNA isolation. To check the quality of RNA, agarose gel electrophoresis was performed. Next, cDNA synthesis of purified RNA was performed using ProtoScript® II First Stand cDNA synthesis Kit (20).

#### RT-qPCR method for Quantitative Efflux pump Gene Expression

Previous studies have demonstrated that heavy metals have a positive effect on the expression of bacterial efflux pumps, which may alter drugs susceptibility towards bacteria (21–24). To quantify the expression of efflux pumps, the *S. aureus* was grown in media with a minimum level of chromium salt or amoxicillin or both. Earlier studies have shown that the β-lactam-related antibiotics are affected by fem factors, which are responsible for peptidoglycan biosynthesis of cell-wall metabolism (19). The norA and mepA are chromosomally encoded efflux pumps of *S. aureus*, which are often used to assess multi-drug resistance profile of *S. aureus* (25–27). Expression levels of femX, mepA and norA was determined using the above-mentioned cDNA and the following PCR primers (Table 2.2 and 2.3) and SYBER-green and PCR-master mix (New England biolabs). To calculate relative gene expression compared to house-keeping GAPDH, comparative threshold cycle was used (25, 28).

**Table 2.1:** Standard clear zone diameter of different antibiotics represents Sensitivity, Moderate Sensitivity and Resistance (CLSI, guideline 2017, M100, 27^th^ Edition.)

| Name of Antibiotic | Disk concentra | Symbol | Zone of Inhibition (diameter in mm) |
| --- | --- | --- | --- |

|  | tion<br>(µg/disk,) |  | Sensitive | Intermediate | Resistance |
| --- | --- | --- | --- | --- | --- |
| Amoxicillin | 10 | AX 10 | ≥29 | - | ≤28 |
| Ampicillin | 10 | AM 10 | ≥27 | - | ≤26 |
| Azithromycin | 15 | AZ 15 | ≥18 | 14-17 | ≤13 |
| Cephadrine | 30 | CE 30 | ≥22 | - | - |
| Chloramphenicol | 30 | C 30 | ≥18 | 13-17 | ≤12 |
| Ciprofloxacin | 5 | CIP 5 | ≥21 | 16-20 | ≤15 |
| Clindamycin | 2 | CD/DA 2 | ≥21 | 15-20 | ≤14 |
| Doxycycline | 30 | DO 30 | ≥16 | 13-15 | ≤12 |
| Erythromycin | 15 | E 15 | ≥23 | 14-22 | ≤13 |
| Methicillin | 5 | MET 5 | ≥18 | - | ≤17 |
| Vancomycin | 30 | VA 30 | ≥15 |  |  |

**Table 2.2:**
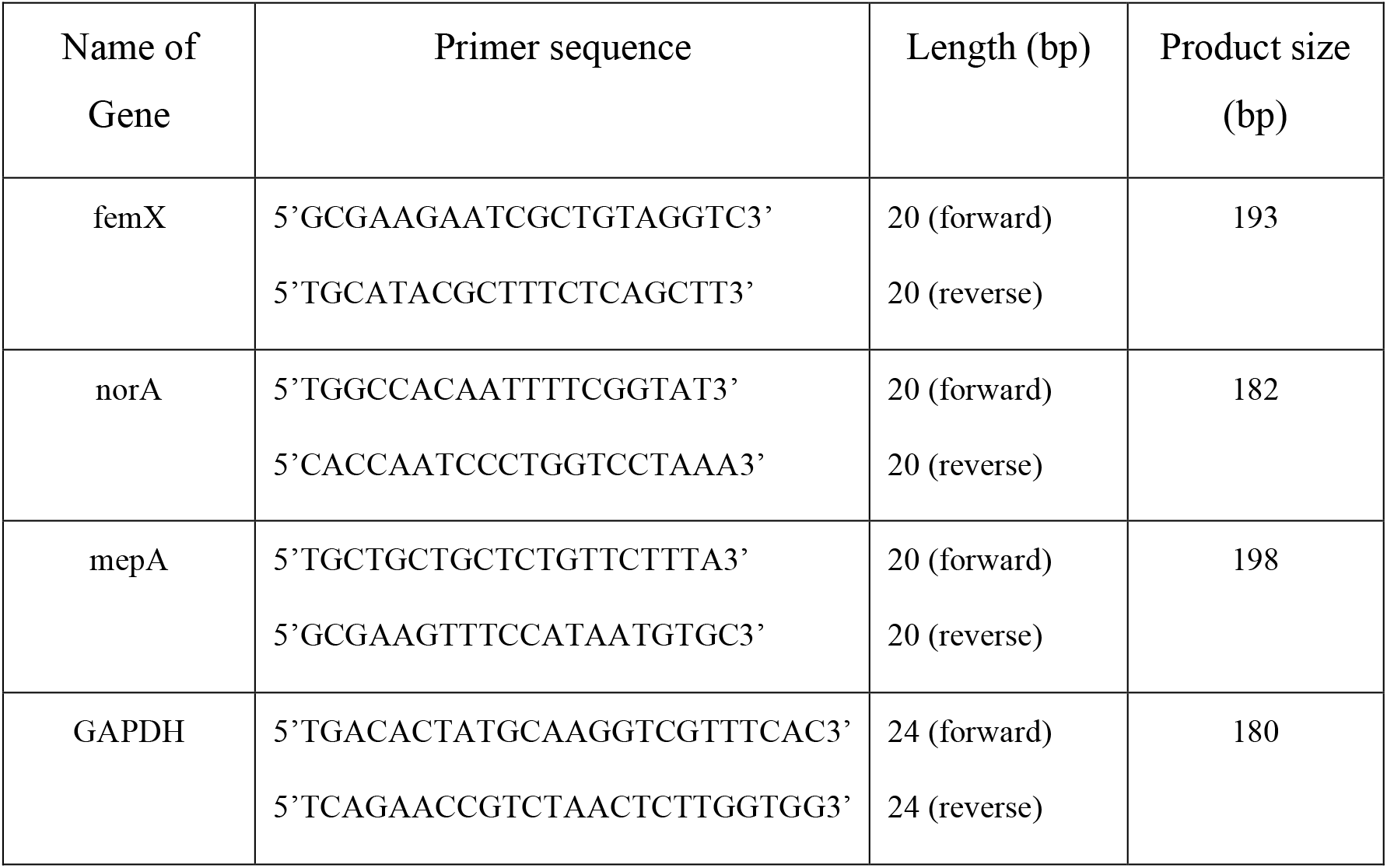
Primer sequence, size and product size

**Table 2.3:** Reagent for RT-qPCR with volumes

| Name of Reagent | Volume |
| --- | --- |
| Template: cDNA | 1.0 μL |
| SYBR-Green master mix | 10 μL |
| 1μM primer Forward (25nM) | 1.0 μL |
| 1μM primer Forward (25nM) | 1.0 μL |
| Nuclease free water | 7.0 µL |
| Total volume | 20 µL |

### Statistical Analysis

All statistical analyses were done using a software called GraphPad Prism version 6.0. To compare the differences of bacterial growth between heavy metal treated and untreated (control) media. The Analysis of Variance (One-way ANOVA) was performed. Data are expressed as mean ± SEM (Standard Error of Mean). Values of p<0.05 are considered as statistically significant.

## 3. Results

### Isolation of *S. aureus* and culture

In this study, *Staphylococcus aureus* ST 80 was isolated from processed raw meat (meat ball) of a local restaurant by Food Microbiology Laboratory, Laboratory Sciences and Services Division, International Centre for Diarrhoeal Disease Research, Bangladesh (ICDDR,B). *S. aureus* is a rod-shaped, Gram-positive and facultative anaerobic microorganism. It is non-fastidious and grow well in Tryptone Soya medium. In this media, at mid-log phase the average viable count of was approximately 1×10^8^ cfu/mL, which was counted from bacterial suspension through Miles-Misra serial dilution method.

### Presence of heavy metals in water bodies or industrial discharge points

In Dhaka, heavy metals are used in the industries like tannery, textile as a source of paints, welding, brazing, soldering, dyes and pigments (29). These industrial effluents are being directly contaminated rivers, cannels and agricultural fields through irrigation channels. Recent studies have shown the presence of heavy metals in the food chains in Bangladesh (4, 30, 31). To measure the levels of heavy metals in the industrial discharge points, we collected water samples from 8 points and measured levels of chromium, cadmium and lead by atomic absorption spectrometry. The concentrations of heavy metals in effluent and river water samples are presented in Table 3.1. The order of heavy metal content is Cr>Pb>Cd with respective concentrations (mg/L) of 2733.10, 0.145 and 0.100 in effluent water which exceeded the WHO (2011) standard.

**Table 3.1:** Presence of chromium, lead and cadmium at industrial wastewater

| Source of Sample | Pb (mg/L) | Cd (mg/L) | Cr (mg/L) |
| --- | --- | --- | --- |
| Tannery-1 (Before ETP) | 0.145 | 0.100 | 2733.10 |
| Tannery-2 (Before ETP) | 0.189 | BDL | 37.54 |
| Tannery-3 (Before ETP) | 2.704 | BDL | 58.54 |
| Textile-2 (Before ETP) | 0.094 | BDL | BDL |
| Textile-1 (Before ETP) | 0.081 | BDL | BDL |
| Textile-1 (After ETP) | 0.140 | BDL | BDL |
| Dhaleshwari River (After ETP) | 1.295 | BDL | 1.52 |
| WHO water quality standard | 0.01 | 0.01 | 0.05 |

Note: ETP-effluent treatment plant. Heavy metals detection limits for Lead 0.04 mg/L, Chromium 0.04 mg/L and Cadmium 0.02 mg/L by Atomic Absorption Spectrophotometer (BDL means below detection level).

### Growth Kinetics of *S. aureus* in the presence of Chromium salt

*Staphylococcus aureus* ST80 was grown in Tryptone soya broth supplemented with 0.5mM, 1mM, 3mM, 5mM, 10mM, 50mM or 100mM chromium salt and in control without chromium. The growth curve (Figure 1A) demonstrated that *Staphylococcus aureus* ST80 tolerates up to 3mM Cr^6+^ salt. The growth kinetics data also showed bacterial growth was inhibited in presence of 5 mM or more concentration of chromium salt. Therefore, 0.5-3.0 mM concentrations of chromium salt were considered as the tolerable level of chromium for *S. aureus* ST80 growth.

**Figure 1:**
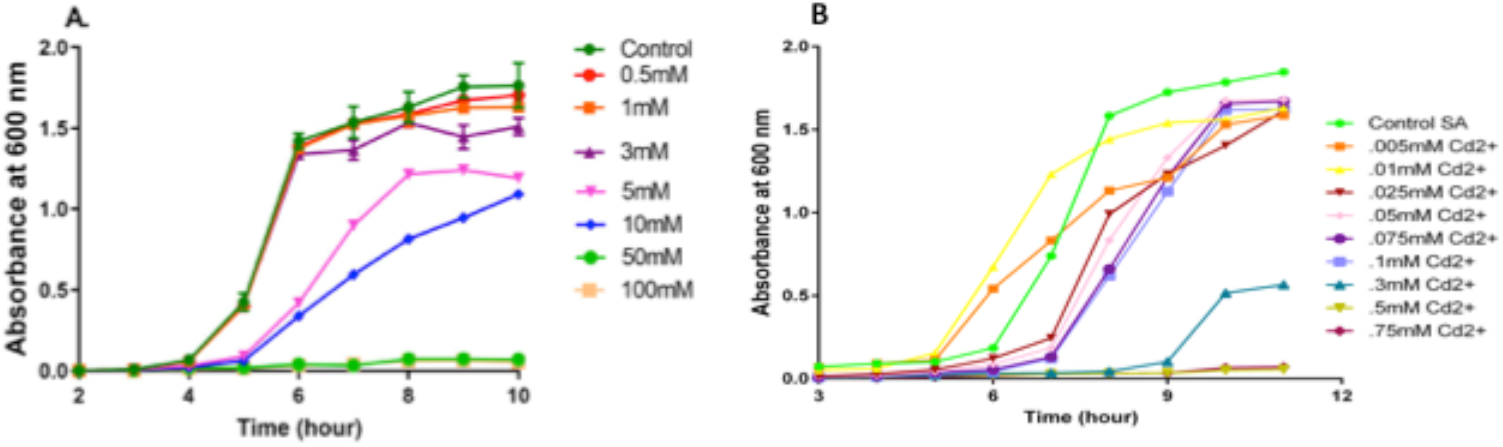
Tolerable levels of chromium and cadmium of *S. aureus* growth. Figure (A) shows the Growth curve of *S. aureus* with different doses of chromium, and figure (B) shows the Growth kinetics of *S. aureus* with different doses of cadmium salt.

On the other hand, *Staphylococcus aureus* ST80 was also grown in Tryptone soya broth in presence of different doses (0.005 mM to 0.75 mM) of cadmium salt. The growth curve (Figure 1B) demonstrated that *Staphylococcus aureus* ST80 tolerates up to 0.1 mM Cd^2+^ salt. The growth kinetics data also showed bacterial growth was inhibited in presence of 0.3 mM or more concentration of cadmium salt. Therefore, 0.005 to 0.1 mM concentrations of cadmium salt were considered as the tolerable level of cadmium for *S. aureus* ST80 growth.

### Screening the pre-exposure effect of chromium or cadmium on antimicrobial sensitivity patterns

A few studies have reported that exposure to metal or heavy metal would not only cause bacteria to develop metal resistance, but antibiotic resistance via co-selection mechanism. However, there is no experimental evidence whether heavy metal like chromium is directly involved to develop multi-drug resistance in *S. aureus*. To examine this hypothesis, Azithromycin 15μg, Chloramphenicol 30μg, Ampicillin 10μg, Amoxicillin 10μg, Erythromycin 15μg, Doxycycline 30μg, Ciprofloxacin 5μg, Cephradine 30μg, Clindamycin 2μg, Methicillin 5μg and Vancomycin 30μg, antibiotic discs were impregnated on Mueller-Hinton or Tryptone Soya agar plates containing chromium or cadmium pre-exposed *S. aureus*. After 20 hours of incubation at 37° C, the antibiotic susceptibility profile was determined by Kirby-Bauer disc diffusion method and the zone of inhibition (ZOI) were interpreted following the Clinical Laboratory Standard Institute (CLSI, USA) guideline, 2017 for *S. aureus*. The antibiotic susceptibility pre-screening data demonstrated that *S. aureus* pre-exposed to either chromium or cadmium, in both conditions showed increased resistance to amoxicillin only compared to heavy metals unexposed control, whereas no significant changes were observed for rest of the antibiotics (Table 3.2).

**Table 3.2:** Screening the pre-exposure effect of chromium or cadmium on antibiotic susceptibility patterns of *Staphylococcus aureus*.

| Name of Antibiotic | Chromium pre-treatment |  |  | Cadmium pre-treatment |  |  |
| --- | --- | --- | --- | --- | --- | --- |
|  | Level of Chromium | ZOI (mm) | p-value | Level of Cadmium | ZOI (mm) | p-value |
| **Amoxicillin** | 0mM<br>0.5mM<br>1.0mM | 36<br>30<br>27 | 0.05 | 0.0<br>0.05mM<br>0.1mM | 37<br>33<br>30 | 0.01 |
| **Ciprofloxacin** | 0mM<br>0.5mM<br>1.0mM | 35<br>34<br>30 | 0.05 | 0.0<br>0.05mM<br>0.1mM | 32<br>30<br>28 | NS |
| **Azithromycin** | 0mM<br>0.5mM<br>1.0mM | 29<br>29<br>26 | NS | 0.0<br>0.05mM<br>0.1mM | 27<br>26<br>26 | NS |
| **Chloramphenicol** | 0mM<br>0.5mM<br>1.0mM | 28<br>27<br>26 | NS | 0.0<br>0.05mM<br>0.1mM | 32<br>31<br>31 | NS |
| **Ampicillin** | 0mM<br>0.5mM<br>1.0mM | 36<br>36<br>33 | 0.05 | 0.0<br>0.05mM<br>0.1mM | 37<br>35<br>35 | NS |
| **Erythromycin** | 0mM<br>0.5mM<br>1.0mM | 30<br>30<br>30 | NS | 0.0<br>0.05mM<br>0.1mM | 32<br>32<br>32 | NS |
| **Clindamycin** | 0mM<br>0.5mM<br>1.0mM | 31<br>31<br>29 | NS | 0.0<br>0.05mM<br>0.1mM | 34<br>33<br>32 | NS |
| **Doxycycline** | 0mM<br>0.5mM<br>1.0mM | 31<br>31<br>29 | 0.05 | 0.0<br>0.05mM<br>0.1mM | 34<br>33<br>31 | NS |
| **Cephadrine** | 0mM<br>0.5mM<br>1.0mM | 32<br>31<br>30 | NS | 0.0<br>0.05mM<br>0.1mM | 32<br>31<br>30 | NS |

Note: NS-non-significance, Data were analyzed The Analysis of Variance (One-way ANOVA) and Sidak multiple comparison tests was performed.

**Table 3.3:**
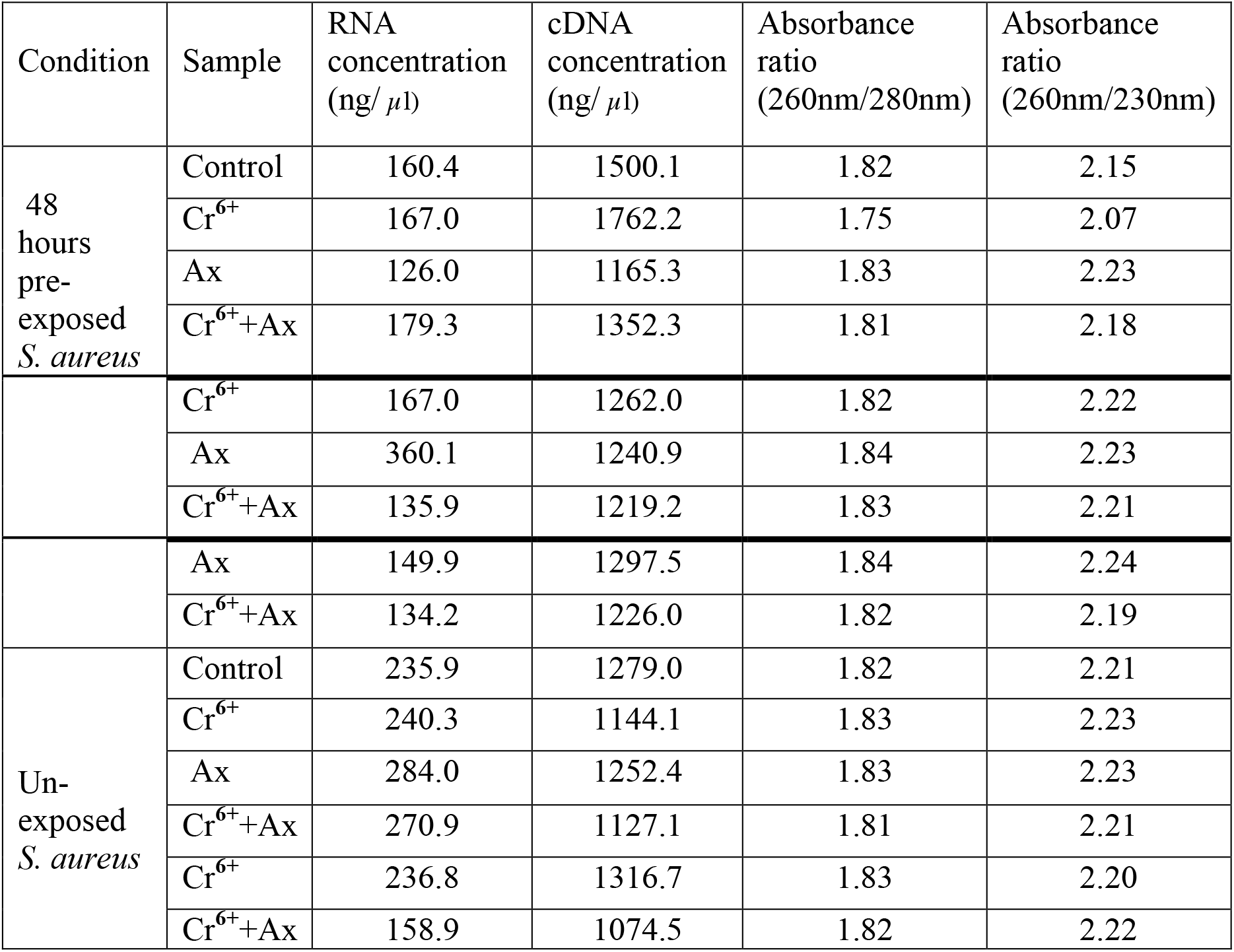
Concentration and purity of RNA of *S. aureus* and the synthesized cDNA.

| Condition | Sample | RNA concentration (ng/ μl) | cDNA concentration (ng/ μl) | Absorbance ratio (260nm/280nm) | Absorbance ratio (260nm/230nm) |
| --- | --- | --- | --- | --- | --- |
| 48 hours pre-exposed *S. aureus* | Control | 160.4 | 1500.1 | 1.82 | 2.15 |
|  | Cr<sup>6+</sup> | 167.0 | 1762.2 | 1.75 | 2.07 |
|  | Ax | 126.0 | 1165.3 | 1.83 | 2.23 |
|  | Cr<sup>6+</sup>+Ax | 179.3 | 1352.3 | 1.81 | 2.18 |
|  | Cr<sup>6+</sup> | 167.0 | 1262.0 | 1.82 | 2.22 |
|  | Ax | 360.1 | 1240.9 | 1.84 | 2.23 |
|  | Cr<sup>6+</sup>+Ax | 135.9 | 1219.2 | 1.83 | 2.21 |
|  | Ax | 149.9 | 1297.5 | 1.84 | 2.24 |
|  | Cr<sup>6+</sup>+Ax | 134.2 | 1226.0 | 1.82 | 2.19 |
| Un-exposed *S. aureus* | Control | 235.9 | 1279.0 | 1.82 | 2.21 |
|  | Cr<sup>6+</sup> | 240.3 | 1144.1 | 1.83 | 2.23 |
|  | Ax | 284.0 | 1252.4 | 1.83 | 2.23 |
|  | Cr<sup>6+</sup>+Ax | 270.9 | 1127.1 | 1.81 | 2.21 |
|  | Cr<sup>6+</sup> | 236.8 | 1316.7 | 1.83 | 2.20 |
|  | Cr<sup>6+</sup>+Ax | 158.9 | 1074.5 | 1.82 | 2.22 |

### Co-exposure Effect of Chromium and Amoxicillin Trihydrate or Cadmium and Amoxicillin on growth of *Staphylococcus aureus*

*S. aureus* ST80 was grown in the Tryptone soya broth, which supplemented with or withour 0.5 mM chromium salt or 0.06 μg/mL Amoxicillin, or both chromium and Amoxicillin. The growth curve (**Figure 2A**) has demonstrated that the chromium pre-exposed *S. aureus* ST80 growth were comparable with chromium un-exposed bacterial control. However, after treated with amoxicillin, the growth of chromium pre-exposed or exposed *S. aureus* was significantly increased compared to chromium un-exposed control. Quantitively, Amoxicillin trihydrate or chromium treatment alone decreased bacterial growth by 52.3% compared to control, whereas the bacterial growth rate was enhanced up to 77.3% in 0.5mM chromium salt and 0.06 μg/mL Amoxicillin trihydrate co-exposed condition. Overall, this growth kinetics data have demonstrated that chromium pre-exposure or re-exposure were minimized the inhibitory effect of Amoxicillin on growth of *S. aureus,* suggesting the existence of chromate enhanced Amoxicillin resistance. Mid-log phase data (**Figure 2B**) demonstrated that bacteria exposed to amoxicillin showed decreased growth compared to control but bacteria pre-exposed to chromium and amoxicillin showed significant changes in growth. Again, bacterial growth under chromium co-exposure and amoxicillin was less than only chromium co-exposed bacteria, but this difficulty was overcome by bacteria pre- and co-exposed to chromium as well as amoxicillin, indicating that amoxicillin, chromium pre- and co-exposed bacteria superceded the growth rate of any single stressed bacteria.

**Figure 2:**
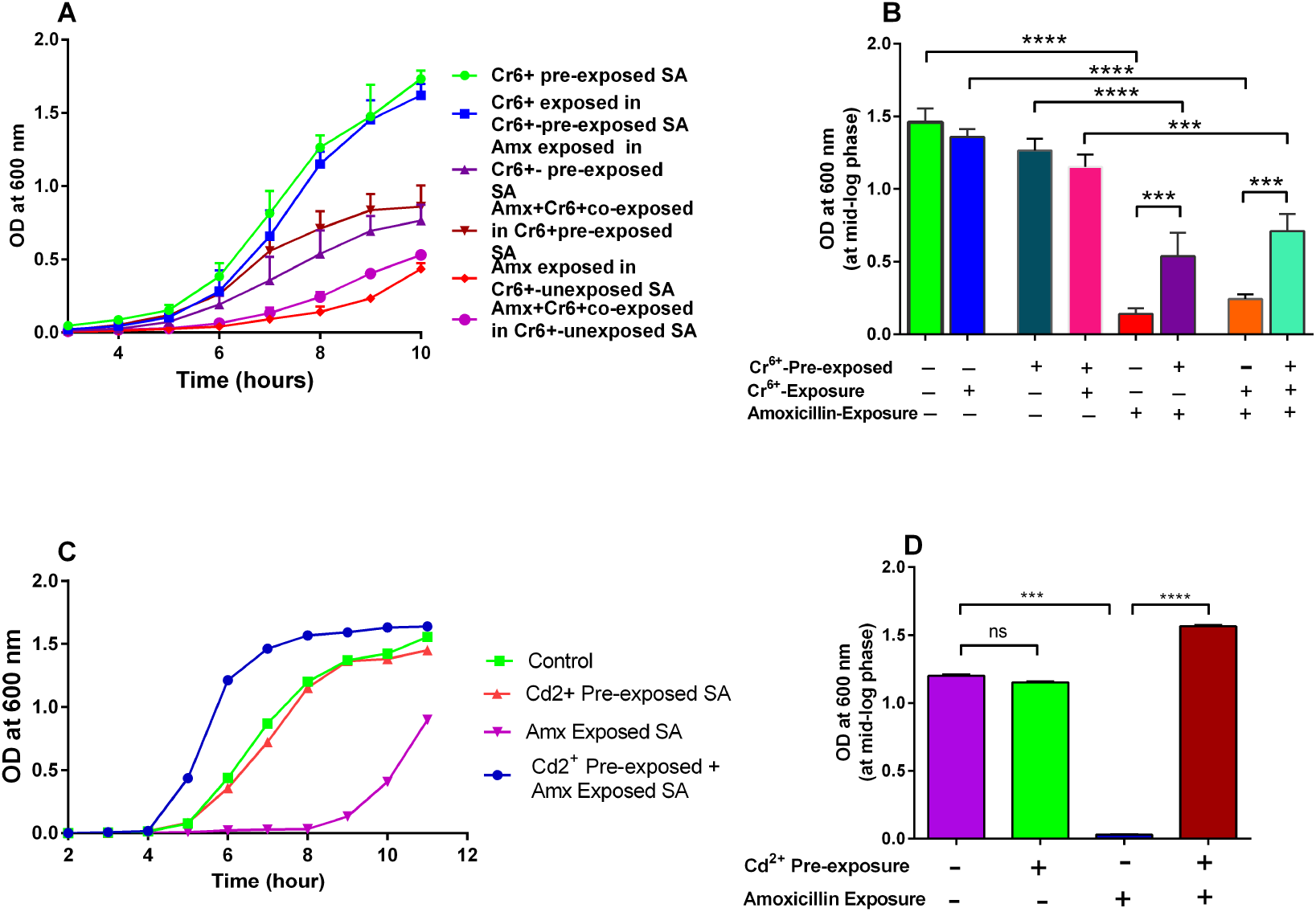
Co-exposure Effect of Chromium and Amoxicillin Trihydrate or Cadmium and Amoxicillin Trihydrate on *Staphylococcus aureus* growth. (A) Growth curve of *Staphylococcus aureus* ST80 in presence of chromium salt and amoxicillin. (B) Growth rate of *Staphylococcus aureus* ST80 at mid-log phase (at 8 hours post-exposure). (C) Growth curve of *Staphylococcus aureus* ST80 in presence of cadmium salt and amoxicillin. (D) Growth rate of *Staphylococcus aureus* ST80 at mid-log phase (at 8 hours post-exposure). 10μl (e.g. 1×10^5^ cfu) of bacterial suspension was added in 30 mL Tryptone Soya broth (TSB) media. The bacterial growth was measured at 600 nm. The Data shows Mean ± SEM and p<0.05, **p<0.01, n=3.

On the other hand, *Staphylococcus aureus* ST80 was also grown in Tryptone soya broth in presence of 0.025 mM cadmium salt or 0.06μg/mL Amoxicillin, or both cadmium and Amoxicillin and in control without any stress. The growth curve (**Figure 2C**) has demonstrated that Amoxicillin trihydrate alone decreased bacterial growth by 52.3% compared to control, whereas this value was enhanced up to 82.6% for cadmium co-exposed bacterial growth rate. That means bacterial growth under both stressed conditions superceded the growth rate of any single stressed bacteria or control. Mid-log phase data (**Figure 2D**) showed that there was no significant changes in terms of growth rate between control and cadmium pre-exposed bacteria, but growth was drastically reduced after amoxicillin treatment. Whereas, Staphylococcus overcame the inhibitory effect of antibiotic by showing increased growth rate in both amoxicillin and pre-exposed cadmium settings.

### Co-exposure effect on the susceptibility of amoxicillin

To determine the minimum inhibitory concentration of Amoxicillin for *S. aureus* under co-exposure of chromium and amoxicillin, the agar dilution method was carried out with Tryptone soya Agar media with or without 0.06 μg/mL to 0.5 μg/mL of Amoxicillin trihydrate. Amoxicillin untreated (e.g. control) agar plate shows plenty of colonies, whereas only 2-3 cfu/per spot were grown in 0.06μg/mL of Amoxicillin trihydrate treated agar plate for untreated bacteria. But no visible bacterial colony was observed in the plates containing 0.125μg/mL of Amoxicillin (**Figure 3A**). Therefore, the minimum inhibitory concentration of Amoxicillin trihydrate for untreated *S. aureus* is 0.06 μg/mL. But for chromium, or Amoxicillin or both treated bacteria, the minimum inhibitory concentration of Amoxicillin trihydrate increased gradually and the data shows that the MIC were 0.125μg/mL for chromium pre-exposed bacteria, 0.25μg/mL for Amoxicillin pre-exposed bacteria and 0.5μg/mL for both chromium and Amoxicillin pre-exposed bacteria. The Antibiotic-Chromium co-exposure assay demonstrated that the MIC for Amoxicillin increased by 8-fold in low-dose of chromium-Amoxicillin co-exposed *S. aureus,* whereas only chromium exposure increased MIC by 2-fold and Amoxicillin exposure increased it by 4-fold compared to control (**Figure 3B**).

**Figure 3:**
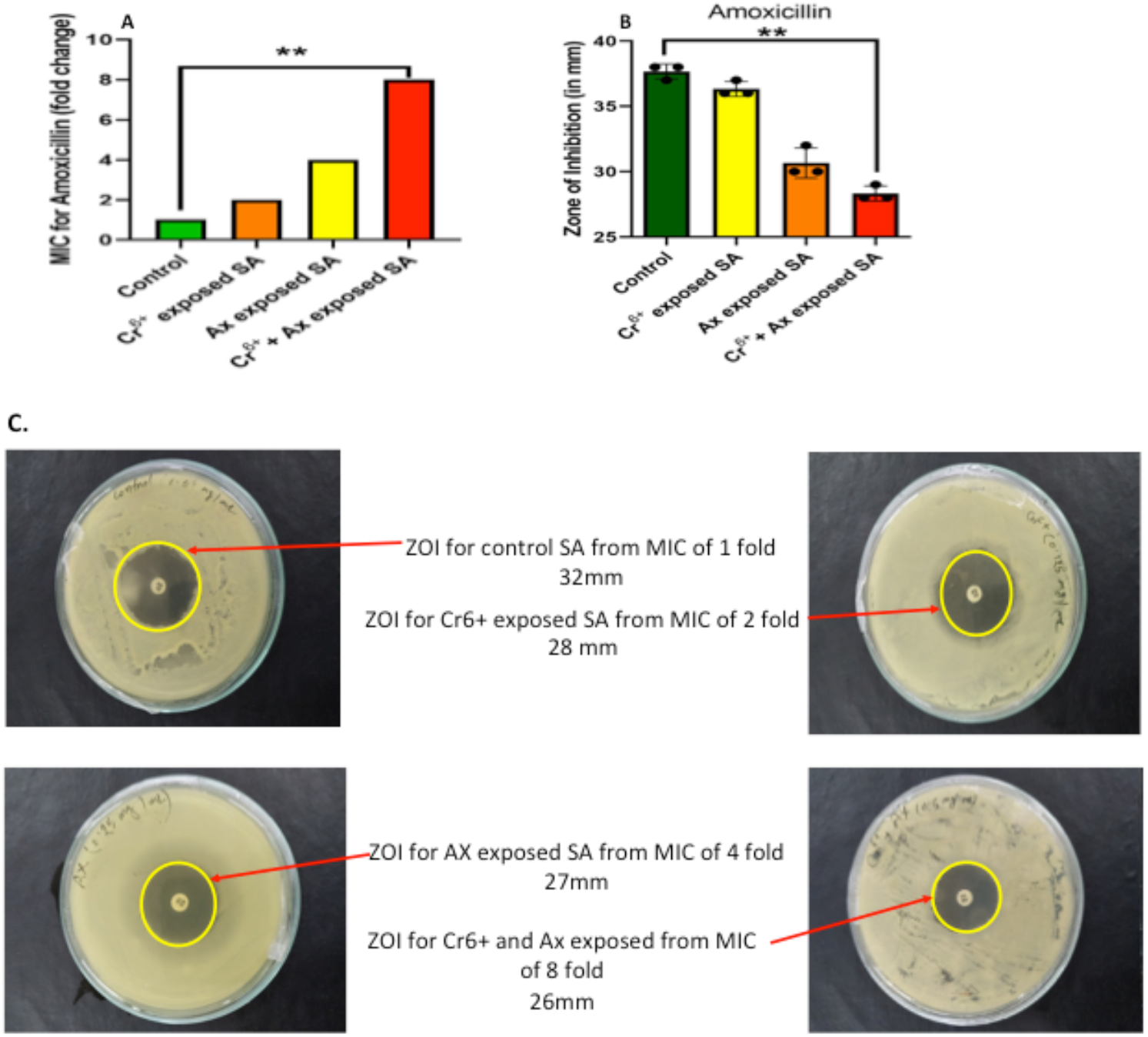
Amoxicillin susceptibility patterns of chromium and amoxicillin pre-exposed *S. aureus:* Figure **(A)** shows the fold change of MIC for amoxicillin. Briefly, *S. aureus* was pre-exposed with a lower dose of chromium and amoxicillin for 48 hours and then 10μL (e.g. 1×10^5^ cfu) of 10 times diluted bacterial suspension was loaded on Tryptone soya Agar plates containing different levels (0.06μg/mL, 0.125μg/mL, 0.25μg/mL, 0.5μg/mL) of Amoxicillin trihydrate and incubated at 37°C for 20 hours. The control media contains no antibiotic. Figure (B) and (C) show the bar chart of ZOI and representative antimicrobial susceptibility test images for amoxicillin. Briefly, *S. aureus* was pre-exposed with a lower dose of chromium and amoxicillin for 48 hours and then 100 μL (e.g. 1×10^7^ cfu) bacterial suspension was spread on each plate and incubated at 37°C for 20 hours. Data are shown as mean + SEM and statistical analysis was performed with one-way ANOVA and Sidak’s post-hoc test, **p<0.01, n=3.

### Antimicrobial susceptibility patterns of chromium pre-exposed *S. aureus*

*Staphylococcus aureus* ST80 was grown in liquid broth media from MIC value of 0.06μg/mL, 0.125μg/mL, 0.25μg/mL, 0.5μg/mL under no stress, only chromium or amoxicillin or both stressed conditions. These bacterial suspensions were spread on Mueller-Hinton Agar Media. From the measurements of zone of inhibition (ZOI) for Amoxicillin antibiotic disc, baseline data shows that *Staphylococcus aureus* ST80 was resistant to Amoxicillin 10μg (e.g. zone of inhibition was 28 mm) under all stressed condition. The ZOI for untreated bacteria was 32 mm, whereas for chromium or amoxicillin or both treatment this value was 28 mm, 27mm and 26 mm, respectively (Figure 3C). Overall, the susceptibility patterns for amoxicillin have demonstrated that co-exposure of a lower dose of chromium and amoxicillin significantly decreased the size of ZOI, which ultimately modify amoxicillin susceptible *S. aureus* to emerge amoxicillin resistant *S. aureus.*

### Co-exposure Effect of cadmium and amoxicillin on susceptibility of amoxicillin

To determine the minimum inhibitory concentration of Amoxicillin for *S. aureus* under co-exposure of cadmium and amoxicillin, the agar dilution method was carried out with Tryptone soya Agar media with or without 0.06μg/mL to 4.0μg/mL of Amoxicillin trihydrate. Amoxicillin untreated (e.g. control) agar plate shows plenty of colonies, whereas only 2-3 cfu/per spot were grown in 0.06μg/mL of Amoxicillin trihydrate treated agar plate for untreated bacteria. But no visible bacterial colonies were observed in the plates containing 0.125μg/mL of Amoxicillin (**Figure 4A**). Therefore, the minimum inhibitory concentration of Amoxicillin trihydrate for untreated *S. aureus* is 0.06μg/mL. But for amoxicillin, cadmium and both treated bacteria the minimum inhibitory concentration (MIC) of Amoxicillin trihydrate was increased gradually and the value is 0.25μg/mL, 1.0 μg/mL, 2.0 μg/mL respectively. The Antibiotic-Cadmium co-exposure assay demonstrated that the MIC for Amoxicillin increased by 32-fold in low-dose of cadmium-Amoxicillin co-exposed *S. aureus,* Whereas, only Amoxicillin exposure increased MIC by 4-fold and cadmium exposure increased it by 16-fold compared to control (**Figure 4B**).

**Figure 4:**
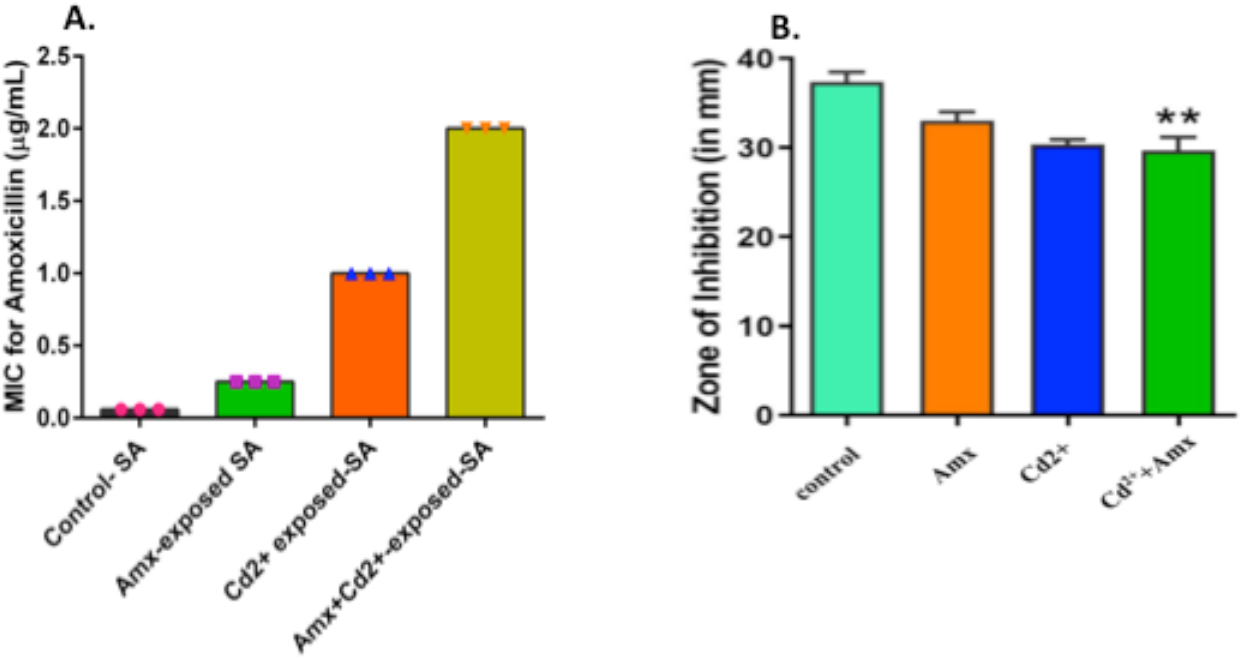
Amoxicillin susceptibility patterns of cadmium and amoxicillin pre-exposed *S. aureus:* Figure (A) shows the MIC for amoxicillin. Briefly, *S. aureus* was pre-exposed with a lower dose of cadmium and amoxicillin for 48 hours and then 10μL (e.g. 1×10^5^ cfu) of 10 times diluted bacterial suspension (adjusted O.D was 0.125 at 600nm) was loaded on Tryptone soya Agar plates containing different levels (0.06μg/mL, 0.125μg/mL, 0.25μg/mL, 0.5μg/mL, 1.0μg/mL, 2.0μg/mL, 4.0μg/mL) of Amoxicillin trihydrate and incubated at 37°C for 20 hours. Figure **(B)** shows the ZOI for Amoxicillin. Briefly, *S. aureus* was pre-exposed with a lower dose of cadmium and amoxicillin for 48 hours and then 100 μL (e.g. 1×10^7^ cfu) bacterial suspension was spread on each plate containing bacteria and incubated at 37°C for 20 hours. Data are shown as mean + SEM and statistical analysis was performed with one-way ANOVA and Sidak’s post-hoc test, **p<0.01, n=3.

### Antibiotic susceptibility pattern of cadmium pre-exposed *S. aureus*

*Staphylococcus aureus* ST80 was further grown in trypsin soya broth medium where the bacterial colonies were inoculated from MIC 1, MIC 4, MIC 16 and MIC 32 of *S. aureus.* These MICs for amoxicillin were emerged due to cadmium and/or amoxicillin pre-exposure in the milieu of S. aureus growth. These bacterial suspensions were spread on Mueller-Hinton Agar Media (Supplementary figure S1). From the measurements of zone of inhibition for Amoxicillin antibiotic disc, baseline data shows that *Staphylococcus aureus* ST80 was resistant to Amoxicillin 10μg (e.g. zone of inhibition was 28 mm) under all stressed condition. The size of zone of inhibition was significantly decreased from untreated to treated condition. Whereas the ZOI for untreated bacteria was 32 mm, for amoxicillin or cadmium or both treatments this value was respectively 27 mm, 26 mm and 25 mm. That means, this resistance patterns for Amoxicillin increased significantly with co-exposure of cadmium and/or amoxicillin (Supplementary figure S2).

### Expression patterns of efflux pumps and Femx gene in *S. aureus*

As demonstrated in antimicrobial susceptibility patterns, co-exposure of amoxicillin and heavy metals like chromium and cadmium alter amoxicillin susceptibility and emerge amoxicillin resistant *Staphylococcus aureus*. We hypothesized that the alteration of antimicrobial susceptibility may be associated with the changes of expression of resistance genes or efflux pumps, which are involved in drug efflux. To address this hypothesis, the expression levels of factor C (femX), efflux pumps mepA and norA of *S. aureus* were assessed by RT-qPCR as RNA level.

The overall quality of an RNA preparation was assessed by electrophoresis on a denaturing agarose gel that would also give some information about RNA yield. A denaturing gel system was used because most RNA form extensive secondary structure via intramolecular base pairing, and thus prevents it from migrating strictly according to its size. **Figure 5A** shows the 5s RNA smear which assures the quality and yield of RNA. Wells with light band denoted disintegrated RNA, which were further extracted. Moreover, to assure the target genes at DNA level and the quality of RT-qPCR gel run was performed. **Figure 5B** indicated the bands for femX, mepA, norA and house-keeping gene GAPDH after RT-qPCR. As the product size of RT-qPCR for the above genes was very close (hence the molecular weight will be very close to each other), the bands of all genes were found in the same alignment after Agarose gel electrophoresis. **Figure 5C** showed that all the targeted genes (femX, mepA, norA, GAPDH) were absolutely amplified following the corresponding C**t** value (number of threshold cycle) for control, amoxicillin, chromium and both amoxicillin-chromium exposed condition. **Figure 5D** denoted that the resistance-related factors *femX, mepA,* and *norA* all were up-regulated in chromium and both chromium-amoxicillin pre-exposed *S. aureus* compared with the control group. *S. aureus* bacterial culture was pre-exposed to chromium salt for 48 hours, which up-regulated the mRNA expression of femX by 25-fold, and mepA by 19-fold and norA by 17-fold compared to control. Whereas, under the co-exposure condition of chromium-amoxicillin, the expression of femX increased by 7-fold, mepA increased by 10-fold and norA increased by 8-fold. In contrast, the expression of these genes was not altered in amoxicillin pre-exposed *S. aureus*. Overall, these findings indicate that heavy metals pre-exposed or heavy metals-antibiotics co-exposed condition up-regulated the expression of drug-resistance genes norA, mepA and femX, which might be responsible for emergence of amoxicillin resistant *S. aureus.*

**Figure 5:**
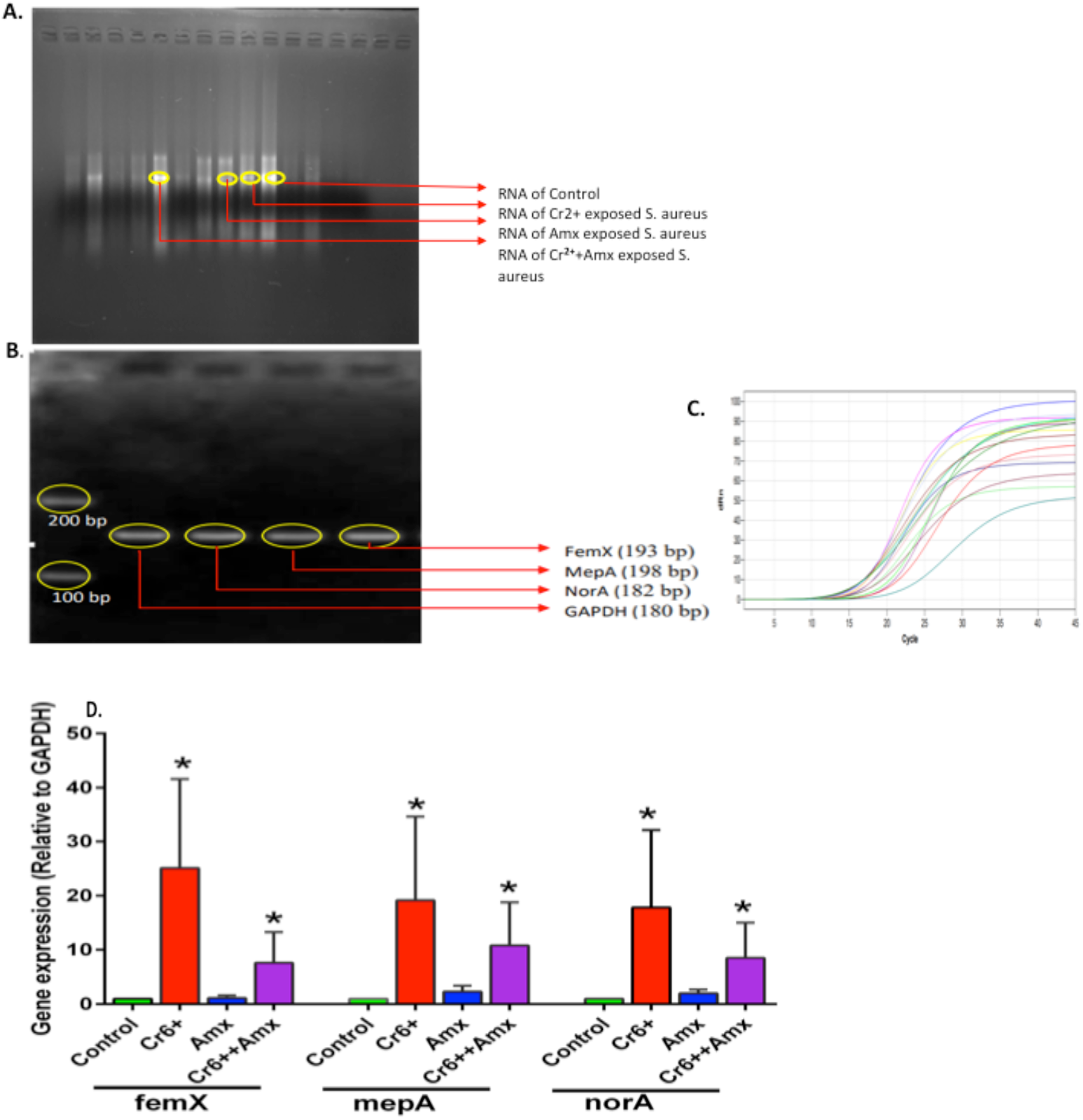
Relative gene expression in chromium and/or amoxicillin pre-exposed *S. aureus*. (A) Band of 5s RNA in Agarose gel electrophoresis. 1.3%, 40 mL gel was prepared with 1X MOPS buffer. Formaldehyde Load Dye was added to each RNA sample as a 1:3 of sample to dye ratio. 8.0 μL sample was loaded on each well. After electrophoresis at 75 V for 30 minutes, the bands were visualized under gel doc. (B) PCR band for product size. (C) Gene amplification. (D) The bar chart for the relative expression of norA, mecA and femx. Data are presented as fold changes on the basis of calculation of gene expression using relative quantification, RQ=2 ^-ΔΔCt^. Data are shown as mean + SEM and statistical analysis was performed with unpaired t-test (two-tailed), non-parametric, Mann whitney t-test. *(n* = 3, asterisk* represents up-regulation, *\*P* < 0.05).

## Discussion

Antimicrobial resistant (AMR) *S. aureus* is one of the leading bacterial causes of infection-associated death, globally (32, 33). *Staphylococci* have acquired AMR-determinants by horizontal gene transfer of mobile genetic elements, by mutations of drug binding sites of target sites and by increasing expression of endogenous efflux pumps (34–36). Industrial processes are responsible for direct deposition of heavy into water, soil, and the atmosphere. Besides, combined contamination of heavy metals and antibiotics contribute to the emergence of multi-drug resistant microbes (2). The deposition of heavy metals in the environment is coupled with their persistence allows for long-term impacts and interactions with microbial communities. The success of bacterial pathogens in the environment is driven by their ability to adapt, spread and establish ecological reservoirs (37). An important determinant of this adaptation is the acquisition of genes that confer resistance, or increase already existing resistance, to antibiotics and heavy metals (37–39).

Studies with Gram-negative bacteria demonstrated that multiple heavy metals/antibiotic resistance genes generally reside on extra chromosomal DNA such as a plasmid (2, 40). One of the main mechanisms for microorganisms’ acquisition of antibiotic resistance under heavy metals selective pressure is i. co-resistance. Favored by evolution, co-resistance, is the occurrence of multiple resistances via the same mobile genetic elements (41). The physical linkage of antibiotic resistance and metal resistance encoded on the plasmid, for example, confers these resistances to the bacteria even when only one co-selecting agent (i.e., antibiotics or heavy metals) is present (40). Whereas, a recent study demonstrated that river isolated *Staphylococci* showed an extra chromosomal independent multiple heavy metals and multiple antibiotic resistance (42). This study also demonstrated plasmid-harboring *Staphylococcus* isolates did not show any effect in their multiple antibiotic and heavy metal resistance profile despite the elimination of all plasmids (42). These observations imply that multi-drug and multi-metal resistant ability of river isolates of Staphylococcus was determined by the chromosome.

In this study, we hypothesized that heavy metals may alter the level of antibiotic susceptibility and facilitate to emerge multi-drug resistant *S. aureus*. To address this hypothesis, the antimicrobial susceptibility profiles were assessed of naturally isolated *Staphylococcus* from processed raw meat and culturally grown *Staphylococcus* with a minimum tolerable level of chromium or cadmium. This investigation shows that *Staphylococcus aureus* ST80 can tolerate up to 3mM K_2_Cr_2_O_7_ and 0.5mM CdCl_2_.H_2_O salts. Previously, a study reported that the minimum inhibition concentrations (MICs) varied from strain to strain of *Staphylococcus (43).* For example the MIC of CuSO_4_.5H_2_O, Cd(NO_3_)_2_.4H_2_O, NaAsO_2_, and ZnSO_4_.7H_2_O was 2, 0.25, 1, 0.25mM for, *Staphylococcus haemolyticus* BB02312, for *Staphylococcus aureus* RN4220 was 4, 0.015, 4, 0.015 mM and for *Staphylococcus haemolyticus* NW19A the MIC was 8, 0.4, 4, 8 mM, respectively (43). Whereas, the MICs of Cu^2+^, Zn^2+^, Cd^2+^, Cr_2_O_7_^2-^, and Ag^+^ respectively were 16, 10, 2.5, 1.6 and 0.25 mM for LSJC7, a member of *Enterobacteriaceae (9).*

The antimicrobial susceptibility profile of this study revealed that minimal inhibitory concentrations of chromium or cadmium and amoxicillin pre-exposure are responsible to emerge amoxicillin resistant *S. aureus*. The alteration of antimicrobial susceptibility in chromium and cadmium salt pre-exposed bacteria might be responsible for emerging antibiotic resistance superbugs in environmental reservoirs. The current observations strongly supported by a recent systemic review, where they showed fourteen published research articles reported the co-occurrence of heavy metal and antibiotic resistance (44). Under selective pressures, microorganisms also acquire antimicrobial resistance by cross-resistance when the route of heavy metals and antimicrobial agents accessed to their target bacteria are similar but not same (2). Evidence of cross-resistance can be found in studies with heavy metal-contaminated environments demonstrating potential microbial adaptation to the environmental selective pressure by acquisition of resistance (40). The most common form of cross-resistance results from the microbial utilization of an efflux pump, a cellular membrane to transport protein (40).

To assess efflux pumps and resistance gene expression at RNA level, the current study adopted the reverse transcription quantitative polymerase chain reaction (RT-qPCR). The RT-qPCR data demonstrated that the expression of femX, mepA and norA have significantly increased in chromium and a lower-dose of amoxicillin pre-exposed *S. aureus* compared to unexposed bacteria. These findings were supported by previous studies with different bacteria, heavy metals and antibiotic co-exposure settings (9, 45, 46). Moreover, previous study also demonstrated that at metabolic stress condition, cell transporters especially efflux pumps, antiporters remove heavy metals and antibiotics from cells in a cross-resistance manner, which is known as cross-regulation (2).

In addition, other studies have reported that the expression of bacterial antibiotic resistance systems were induced by heavy metals, for example, the transcription of multi-drug efflux pump genes acrD and mdtABC in *Salmonella enterica,* were induced by a two-component signal transduction system BaeRS in response to copper or zinc resulting in enhanced antibiotic resistance (45). Besides, chromate or copper modifies SoxS regulator and hence enhances the expression of multi-drug efflux pump AcrAB-TolC in *E. coli* (47). These proposed efflux pumps (e.g. AcrAB-TolC) are mostly responsible for conferring resistance against diverse antibiotics (47). Recently, in a tissue cage infection model study demonstrated that in-complete or improper regimen of amoxicillin against *S. aureus* cause to induce amoxicillin resistance genes such as mecA, femA, femB and femx gene (46). Another study has demonstrated that heavy metals with no nutritional benefits for microorganisms cause oxidative stress (48) that might trigger adaptations and bacterial growth in heavy metals and antimicrobial co-contamination settings. Presumably, high and chronic exposure to heavy metals and metals cause irreversible damage to bacterial DNA and the cell membrane, which later acts as an environmental selector for cellular defense (49). This defense pathway could habituate bacteria to grow in presence of heavy metals, which may contribute to acquire resistance to heavy metals and antibiotics as a cross-resistance fashion. The recent study on bacterial resistance to heavy metals supported these earlier observations, where they showed that whether an intrinsic, natural, or a selective pressure-induced modification, bacteria acquire antimicrobial resistance through increases co- and/or cross-resistance pathways (50). These observations and current study findings have strongly supported a mechanistic explanation (e.g. cross-resistance/regulation) behind the emergence of amoxicillin resistant *S. aureus* in the heavy metal and antibiotic co-contamination setting.

### Conclusion

This study demonstrates culture-based and molecular-based methods for chromium or cadmium and a lower-dose of amoxicillin pre-exposure is responsible to emerge amoxicillin resistant *S. aureus*. However, it is warranted to validate the current observed antibiotic resistant patterns in the corresponding efflux pumps knockout model of *S. aureus.* The efflux pump may effectively regulate the interaction of both heavy metals and antibiotics by conferring resistance.

## ACKNOWLEDGMENTS

The work is supported by two research grants funded by the Ministry of Education, Bangladesh (Banbeis, GARE ID LS2019989) and the Ministry of Science and Technology, Bangladesh (2021-2022). The Atomic Absorption Spectroscopy is supported by Bangladesh Council of Scientific and Industrial Research (BCSIR). RNA extraction, RT-qPCR and Biosafety-2 supported by Molecular Biology lab Genomic research lab and BSL-2 lab, department of Biochemistry and Molecular Biology, University of Dhaka, Bangladesh.

## AUTHOR CONTRIBUTIONS

TNI and FSM have equally conducted and analysed experiments. RY optimized RT-qPCR with bacterial RNA. MBI isolated bacterial strain. TR and DHD reviewed the manuscript. MM designed experiments and reviewed analysis with TNI and FSM. MM, TNI and FSM wrote the manuscript with input of all authors.

## REFERENCES

1. Zhang R, Eggleston K, Rotimi V, Zeckhauser RJ. Antibiotic resistance as a global threat: evidence from China, Kuwait and the United States. Global Health. 2006;2(1):6.

2. Baker-Austin C, Wright M, Stepanauskas R, McArthur J. Co-selection of antibiotic and metal resistance. Trends in Microbiology. 2006;14(4):176–82.

3. Seiler C, Berendonk T. Heavy metal driven co-selection of antibiotic resistance in soil and water bodies impacted by agriculture and aquaculture. Frontiers in Microbiology. 2012;3.

4. Ahmad JU, Goni MA. Heavy metal contamination in water, soil, and vegetables of the industrial areas in Dhaka, Bangladesh. Environ Monit Assess. 2010;166(1-4):347–57.

5. Gupta N, Khan DK, Santra SC. An assessment of heavy metal contamination in vegetables grown in wastewater-irrigated areas of Titagarh, West Bengal, India. Bull Environ Contam Toxicol. 2008;80(2):115–8.

6. Martínez JL, Rojo F. Metabolic regulation of antibiotic resistance. FEMS Microbiol Rev. 2011;35(5):768–89.

7. Zhu Y-G, Johnson TA, Su J-Q, Qiao M, Guo G-X, Stedtfeld RD, et al. Diverse and abundant antibiotic resistance genes in Chinese swine farms. Proc Natl Acad Sci U S A. 2013;110(9):3435–40.

8. Looft T, Johnson TA, Allen HK, Bayles DO, Alt DP, Stedtfeld RD, et al. In-feed antibiotic effects on the swine intestinal microbiome. Proc Natl Acad Sci U S A. 2012;109(5):1691–6.

9. Chen S, Li X, Sun G, Zhang Y, Su J, Ye J. Heavy metal induced antibiotic resistance in bacterium LSJC7. Int J Mol Sci. 2015;16(10):23390–404.

10. Heuer H, Schmitt H, Smalla K. Antibiotic resistance gene spread due to manure application on agricultural fields. Curr Opin Microbiol. 2011;14(3):236–43.

11. Khan S, Cao Q, Zheng YM, Huang YZ, Zhu YG. Health risks of heavy metals in contaminated soils and food crops irrigated with wastewater in Beijing, China. Environ Pollut. 2008;152(3):686–92.

12. Knapp CW, McCluskey SM, Singh BK, Campbell CD, Hudson G, Graham DW. Antibiotic resistance gene abundances correlate with metal and geochemical conditions in archived Scottish soils. PLoS One. 2011;6(11):e27300.

13. Peltier E, Vincent J, Finn C, Graham DW. Zinc-induced antibiotic resistance in activated sludge bioreactors. Water Res. 2010;44(13):3829–36.

14. Berg J, Tom-Petersen A, Nybroe O. Copper amendment of agricultural soil selects for bacterial antibiotic resistance in the field. Lett Appl Microbiol. 2005;40(2):146–51.

15. Lowy FD. Staphylococcus aureus infections. New England journal of medicine. 1998;339(8):520–32.

16. Ahsan MA, Satter F, Siddique MAB, Akbor MA, Ahmed S, Shajahan M, et al. Chemical and physicochemical characterization of effluents from the tanning and textile industries in Bangladesh with multivariate statistical approach. Environ Monit Assess. 2019;191(9):575.

17. Hassan M, Rahman MATMT, Saha B, Kamal AKI. Status of heavy metals in water and sediment of the Meghna river, Bangladesh. Am J Environ Sci. 2015;11(6):427–39.

18. Wiegand I, Hilpert K, Hancock REW. Agar and broth dilution methods to determine the minimal inhibitory concentration (MIC) of antimicrobial substances. Nat Protoc. 2008;3(2):163–75.

19. Bera A, Biswas R, Herbert S, Kulauzovic E, Weidenmaier C, Peschel A, et al. Influence of wall teichoic acid on lysozyme resistance in Staphylococcus aureus. J Bacteriol. 2007;189(1):280–3.

20. Atshan SS, Shamsudin MN, Lung LTT, Ling KH, Sekawi Z, Pei CP, et al. Improved method for the isolation of RNA from bacteria refractory to disruption, including S. aureus producing biofilm. Gene. 2012;494(2):219–24.

21. Blair JMA, Webber MA, Baylay AJ, Ogbolu DO, Piddock LJV. Molecular mechanisms of antibiotic resistance. Nat Rev Microbiol. 2015;13(1):42–51.

22. Karam G, Chastre J, Wilcox MH, Vincent J-L. Antibiotic strategies in the era of multidrug resistance. Crit Care. 2016;20(1):136.

23. Spengler G, Kincses A, Gajdács M, Amaral L. New Roads leading to old destinations: Efflux pumps as targets to reverse multidrug resistance in bacteria. Molecules. 2017;22(3):468.

24. Webber MA, Piddock LJV. The importance of efflux pumps in bacterial antibiotic resistance. J Antimicrob Chemother. 2003;51(1):9–11.

25. Huet AA, Raygada JL, Mendiratta K, Seo SM, Kaatz GW. Multidrug efflux pump overexpression in Staphylococcus aureus after single and multiple in vitro exposures to biocides and dyes. Microbiology. 2008;154(Pt 10):3144–53.

26. Kaatz GW, DeMarco CE, Seo SM. MepR, a repressor of the Staphylococcus aureus MATE family multidrug efflux pump MepA, is a substrate-responsive regulatory protein. Antimicrob Agents Chemother. 2006;50(4):1276–81.

27. Paulsen IT, Brown MH, Skurray RA. Proton-dependent multidrug efflux systems. Microbiol Rev. 1996;60(4):575–608.

28. Livak KJ, Schmittgen TD. Analysis of relative gene expression data using real-time quantitative PCR and the 2(-Delta Delta C(T)) Method. Methods. 2001;25(4):402–8.

29. Wu X, Cobbina S, Mao G, Xu H, Zhang Z, Yang L. A review of toxicity and mechanisms of individual and mixtures of heavy metals in the environment. Environmental Science and Pollution Research. 2016;23(9):8244–59.

30. Ahmad JU GM. Heavy metal contamination in water, soil, and vegetables of the industrial areas in Dhaka, Bangladesh 2010; 166:[347–57 pp.].

31. Ahmad MK IS, Rahman S, Haque M, Islam MM. Heavy metals in water, sediment and some fshes of Buriganga River, Bangladesh 2010; 4(2):[321–32 pp.].

32. Collaborators GAR. Global mortality associated with 33 bacterial pathogens in 2019: a systematic analysis for the Global Burden of Disease Study 2019. Lancet. 2022;400(10369):2221–48.

33. Gona PN, More AF. Bacterial pathogens and climate change. Lancet. 2022;400(10369):2161–3.

34. Jensen SO, Lyon BR. Genetics of antimicrobial resistance in Staphylococcus aureus. Future Microbiol. 2009;4(5):565–82.

35. Foster TJ. Antibiotic resistance in Staphylococcus aureus. Current status and future prospects. FEMS Microbiol Rev. 2017;41(3):430–49.

36. Pennone V, Prieto M, Álvarez-Ordóñez A, Cobo-Diaz JF. Antimicrobial Resistance Genes Analysis of Publicly Available. Antibiotics (Basel). 2022;11(11).

37. Hanssen AM, Ericson Sollid JU. SCCmec in staphylococci: genes on the move. FEMS Immunol Med Microbiol. 2006;46(1):8–20.

38. Aktan Y, Tan S, Icgen B. Characterization of lead-resistant river isolate Enterococcus faecalis and assessment of its multiple metal and antibiotic resistance. Environ Monit Assess. 2013;185(6):5285–93.

39. Ozer G, Ergene A, Icgen B. Biochemical and Molecular Characterization of Strontium-resistant Environmental Isolates of Pseudomonas fluorescens and Sphingomonas paucimobilis. Geomicrobiology Journal. 2013;30(5):381–90.

40. Pal C, Bengtsson-Palme J, Kristiansson E, Larsson DG. Co-occurrence of resistance genes to antibiotics, biocides and metals reveals novel insights into their co-selection potential. BMC Genomics. 2015;16:964.

41. Ye J, Rensing C, Su J, Zhu YG. From chemical mixtures to antibiotic resistance. J Environ Sci (China). 2017;62:138–44.

42. Yilmaz F, Orman N, Serim G, Kochan C, Ergene A, Icgen B. Surface water-borne multidrug and heavy metal-resistant Staphylococcus isolates characterized by 16S rDNA sequencing. Bull Environ Contam Toxicol. 2013;91(6):697–703.

43. Xue H, Wu Z, Li L, Li F, Wang Y, Zhao X. Coexistence of Heavy Metal and Antibiotic Resistance within a Novel Composite Staphylococcal Cassette Chromosome in a Staphylococcus haemolyticus Isolate from Bovine Mastitis Milk. Antimicrobial Agents and Chemotherapy. 2015;59(9):5788–92.

44. Nguyen CC, Hugie, C.N., Kile, M.L. et al. Association between heavy metals and antibiotic-resistant human pathogens in environmental reservoirs: A review 2019; 46(13).

45. Nishino K, Nikaido E, Yamaguchi A. Regulation of multidrug efflux systems involved in multidrug and metal resistance of Salmonella enterica serovar Typhimurium. J Bacteriol. 2007;189(24):9066–75.

46. Yao Q, Gao L, Xu T, Chen Y, Yang X, Han M, et al. Amoxicillin Administration Regimen and Resistance Mechanisms of. Front Microbiol. 2019;10:1638.

47. Harrison JJ, Tremaroli V, Stan MA, Chan CS, Vacchi-Suzzi C, Heyne BJ, et al. Chromosomal antioxidant genes have metal ion-specific roles as determinants of bacterial metal tolerance. Environ Microbiol. 2009;11(10):2491–509.

48. T K. Online textbook of bacteriology 2012.

49. Safari Sinegani AA, Younessi N. Antibiotic resistance of bacteria isolated from heavy metal-polluted soils with different land uses. J Glob Antimicrob Resist. 2017;10:247–55.

50. Gorovtsov AV, Sazykin IS, Sazykina MA. The influence of heavy metals, polyaromatic hydrocarbons, and polychlorinated biphenyls pollution on the development of antibiotic resistance in soils. Environ Sci Pollut Res Int. 2018;25(10):9283–92.

