## Supplementary data for "Chromium/cadmium plays a pivotal role to emerge amoxicillin resistant *Staphylococcus aureus*"

**Legend S1: Determination of Minimum inhibitory concentration of Amoxicillin trihydrate by Agar Dilution Method.** 10µL (e.g. $1x10^5$ cfu) of 10 times diluted bacterial suspension (adjusted O.D was 0.125 at 600nm) was spotted on Tryptone soya Agar plates containing different levels (0.06µg/mL, 0.125µg/mL, 0.25µg/mL, 0.5µg/mL) of Amoxicillin trihydrate and incubated at $37^0C$ for 20 hours.

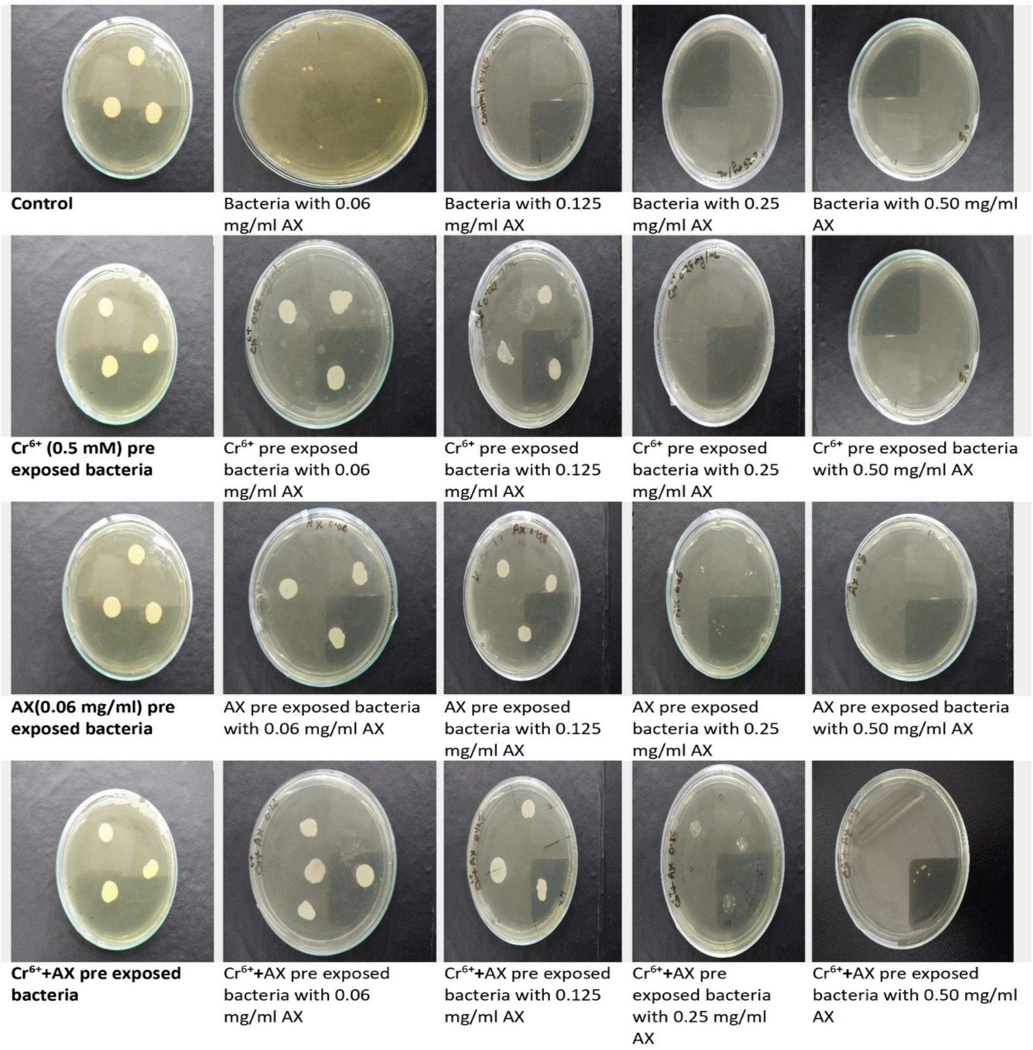


Figure S1 Determination of Minimum inhibitory concentration of Amoxicillin trihydrate by Agar Dilution Method

Legend S2: **Determination of Minimum inhibitory concentration of Amoxicillin trihydrate by Agar Dilution Method.** 10µL (e.g. $1x10^5$ cfu) of 10 times diluted bacterial suspension (adjusted O.D was 0.125 at 600nm) was spotted on Tryptone soya Agar plates containing different levels (0.06µg/mL, 0.125µg/mL, 0.25µg/mL, 0.5µg/mL,1.0µg/mL,2.0µg/mL,4.0µg/mL) of Amoxicillin trihydrate and incubated at $37^0C$ for 20 hours.

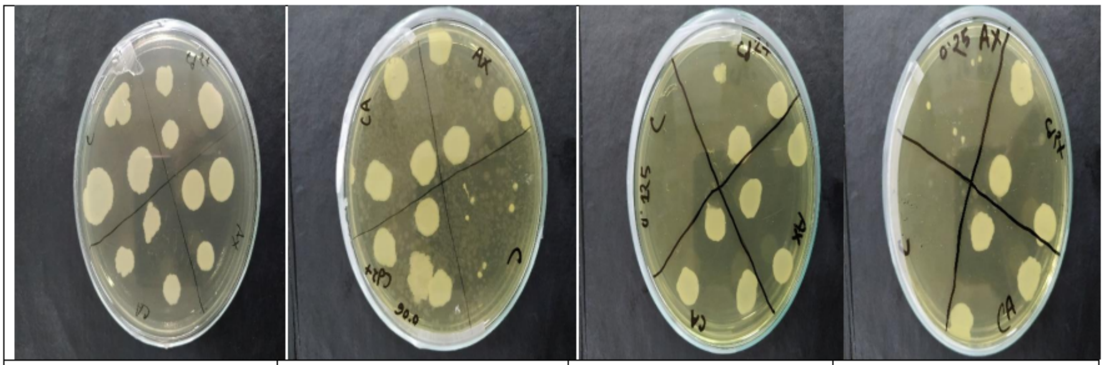

| | | | |
|---|---|---|---|
| Stressed(0.06 mg/ml AX,0.025mM $Cd^{2+}$,both exposure) and unstressed bacteria with no antibiotic | Stressed(0.06 mg/ml AX,0.025mM $Cd^{2+}$,both exposure) and unstressed bacteria with 0.06 mg/ml AX | Stressed(0.06 mg/ml AX,0.025mM $Cd^{2+}$,both exposure) and unstressed bacteria with 0.125 mg/ml AX | Stressed(0.06 mg/ml AX,0.025mM $Cd^{2+}$,both exposure) and unstressed bacteria with 0.25 mg/ml AX |

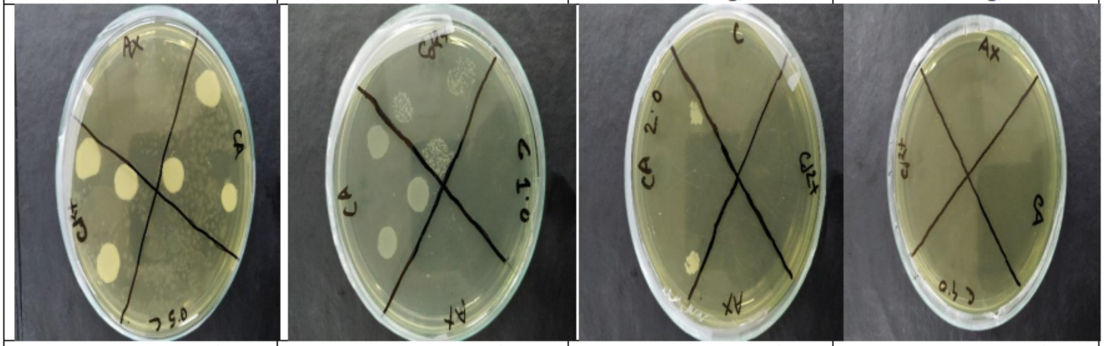

| | | | |
|---|---|---|---|
| Stressed(0.06 mg/ml AX,0.025mM $Cd^{2+}$,both exposure) and unstressed bacteria with 0.5 mg/ml AX | Stressed(0.06 mg/ml AX,0.025mM $Cd^{2+}$,both exposure) and unstressed bacteria with 1.0 mg/ml AX | Stressed(0.06 mg/ml AX,0.025mM $Cd^{2+}$,both exposure) and unstressed bacteria with 2.0 mg/ml AX | Stressed(0.06 mg/ml AX,0.025mM $Cd^{2+}$,both exposure) and unstressed bacteria with 4.0 mg/ml AX |

Figure S2 **Determination of Minimum inhibitory concentration of Amoxicillin trihydrate by Agar Dilution Method.**
